# Establishment and Characterization of Three Novel CAF Cell Lines from HNSCC Patients

**DOI:** 10.1101/2023.01.08.523131

**Authors:** Nehanjali Dwivedi, Hafsa Bahaar, DN Shashank, Christine Elizabeth Cherry, Charitha Gangadharan, Amritha Suresh, Moni A Kuriakose, Vijay Pillai, Manjula Das, P.K Smitha

## Abstract

Due to the high rates of tobacco chewers, smokers, and alcohol consumers in India, Head and Neck Squamous Cell Carcinoma (HNSCC) is one of the primary causes of mortality. Being profoundly varied in nature, treating patients diagnosed with HNSCC can be difficult. An in vitro cell line model is needed to better comprehend the heterogeneity especially via the interaction with components of the microenvironment like Cancer Associated Fibroblasts (CAFs). The effectiveness of creating cell lines from head and neck cancers is, however, poor. Furthermore, except for the two reported earlier by us, no other CAF cell lines are available to study the cross-talk of the tumor with its microenvironment. In this study, we report three novel CAF lines, spontaneously immortalized from HPV negative male patients with habits of tobacco and diagnosed with squamous cell carcinoma of the upper alveolus, larynx and buccal mucosa as opposed to HPV positive non-habitual female patients with cancer of the tongue as previously reported. The CAFs increased the tumorigenicity of epithelial cells in indirect co-culture experiments. Negative staining with EpCAM, CD31 and CD45, while positive staining with FSP-1 determined their fibroblast specific lineage. The developed CAF cultures are the first of their kind from the mentioned sites, and they will be an invaluable tool for learning how tumor-stroma and tumor interact with one another and discovering newer targets and pathways for treating HNSCC.

## Introduction

Head and Neck Squamous Cell Carcinoma (HNSCC), the seventh most common cancer worldwide accounts for more than 325,000 mortalities and 660,000 new incidences across the world (1,2). HNSCC is the most prevalent form of male cancer in India, which may be related to the population’s extensive exposure to risky lifestyle parameters such as tobacco, cigarette smoking, areca nut chewing, and drinking (3,4). Infection with the human papilloma virus (HPV) has recently been linked to an increased risk of HNSCC (5–7). In India, the 5-year survival rates for patients with HNSCC at the early and advanced stages are 82 and 27%, respectively (8). The five-year survival rate of patients with HNSCC is still low due to distant metastasis and recurrence, and the patient’s quality of life has remained a severe issue despite advances in early detection and therapy (4,9). Therefore, a deeper comprehension of the intricate etiology of HNSCC is required to create a more potent treatment.

An essential part of the growth of tumors is the extracellular matrix (ECM), the macromolecule-rich tumor microenvironment (TME). It is now understood that cell interactions within the tumor microenvironment play a key role in tumor development and progression (10). Fibroblasts are the second most common type of cells in the microenvironment and are a dynamic population of cells with a wide range of phenotypic and functional characteristics. Nearly all solid tumor tissues include CAFs, which are crucial for the malignant development of cancer, including metastasis and the epithelial-to-mesenchymal transition (EMT) (11). CAFs, therefore can be viewed as “the other side of the coin” in tumorigenesis (12).

Cell lines remain a valuable resource for performing translational research in laboratory settings despite the challenges of long-term preservation (13). Establishment of primary cultures enables investigation into the parent tumor’s microenvironment-driven traits. Additionally, it is imperative to create and characterize novel CAF lines from patient malignancies due to the dearth of commercially available CAFs.

A key tool for improving our understanding of tumor heterogeneity and stroma-tumor cross-talk is in vitro cell-based models of cancer (14–16). Tatake et.al, discuss the development of epithelial cell lines from the lower alveolus, buccal mucosa, gingivobuccal mucosa, oral cavity, tongue, sinonasal, pyriform, and retromolar trigone in the treatment of head and neck cancer (17). 85 cell lines from distinct head and neck locations were further assembled and described by Zhao et al. (18) in 2011. Hayes et al. previously produced 16 cell lines from patients with HNSCC to better understand the effects of various mutations on tumor behavior (19). However, establishment of CAF cell lines, so far, have only been reported from this group in June 2022 (20).

The aim of the present study was, therefore, to establish novel in-vitro cell-based study platforms in the form of CAFs from patients diagnosed with moderately differentiated squamous cell carcinoma of the larynx, upper alveolus and buccal mucosa. Unlike our previous report, all the patients reported in this study were males and had smoking, chewing and drinking habits and were HPV negative. Studying the effect and cross-talk of these CAFs with the epithelial cell lines from oral cancer can therefore be an essential tool in understanding head and neck cancer progression.

## Materials and Methods

### Tumor sample collection and cell line establishment

Tumor samples were collected from three different patients diagnosed with moderately differentiated squamous cell carcinoma, after receiving the patient’s informed consent form. The current study was approved by the Narayana Health City ethics committee with an approval number – NHH/MECCL2015405 (A) from Bengaluru, India. Tumor samples were collected in sterile RPMI-1640 (cat.no. AT222A; HiMedia Laboratories; LLC) supplemented with 3X Penicillin-Streptomycin solution (cat. no. 15140122; Gibco; Thermo Fisher Scientific, Inc.) from two males with habits of drinking, chewing and smoking, while for one of the patients, the habits were not reported. Patients MhCL03-F, MhCA04-F, and MhCB05-F were diagnosed with squamous cell carcinoma of the larynx, upper alveolus, and buccal mucosa respectively. While patients MhCL03-F and MhCA04-F presented with primary tumor, patient MhCB05-F was from a recurrent patient. The clinical details of the patients are mentioned in Table I. Tumor samples (5mm punch) were digested with trypsin for MhCB05-F patient and collagen for MhCL03-F and MhCA04-F patients for 15 min at 37□ after decontamination with povidone-iodine solution (Win Medicare Pvt. Ltd.). The tissue was then chopped into smaller sections and placed in serrated petri dishes in RPMI-1640 medium supplemented with 20% FBS (cat.no. RM10434; HiMedia Laboratories; LLC) and 1X Penicillin-Streptomycin solution and incubated at 37□ with 5% CO_2_. Dead cells and debris were removed by changing the culture medium every 48 h. The resulting fibroblast cells were then trypsinized and passaged for more than 40 passages in RPMI complete medium and characterized further. The names have been arrived via an acronym, where, M – Mazumdar shaw medical foundation; h – human; C – Cancer of; L – Larynx/A – upper Alveolus/B – Buccal mucosa; 03/04/05 – Patient code; F/E – Fibroblast/Epithelial.

**Table I.** Clinical and pathological details of the established cell lines.

| Cell line | Patient Age (years)/sex/habit | Tumor diagnosis | Pathologic al Staging | Comorbidity | Tumor Progression |
| --- | --- | --- | --- | --- | --- |
| MhCL03-F | 65/M/Smoking | Moderately differentiated Squamous cell carcinoma of the Larynx | T4bN2bMx | Diabetes | Presentation: Primary |
| MhCA04-F | 80/M/Drinking and Chewing | Moderately differentiated Squamous cell carcinoma of the upper alveoulus | T4 Nx Mx. | Diabetes / HTN/ Angiogram | Presentation : Primary |
| MhCB05-F | 65/M/Not reported | Moderately differentiated Squamous cell carcinoma of the buccal mucosa | T4bN0 | Asmatic | Presentation: Primary tumor in jan 2016 treated by surgerty and no adjuvant therapy. Recurrence in jan 2017 |

M: Male; HTN: Hypertension

### HeLa cell culture

HeLa cells were grown in DMEM medium (cat. no. 11995-065; Gibco; Thermo Fisher Scientific, Inc.), pH 7.2, supplemented with 10% FBS and 1X penicillin-streptomycin solution at 37 °C with 5% CO_2_. The authenticity of HeLa cell line was further confirmed by STR profiling (Table II).

**Table II.** STR analysis of HeLa cell line. The STR profile matched 100% to HeLa cell line from American Type Culture Collection (ATCC) database (21) as authenticated by TheraCUES.

|  | Marker | Allele#1 | Allele#2 |
| --- | --- | --- | --- |
| HeLa | TH01 | 7 | 7 |
|  | D21S11 | 27 | 28 |
|  | D5S818 | 11 | 12 |
|  | D13S317 | - | - |
|  | D7S820 | 8 | 12 |
|  | D16S539 | 9 | 10 |
|  | CSF1PO | 9 | 10 |
|  | AMEL | X | X |
|  | vWA | 16 | 18 |
|  | TPOX | 8 | 12 |

### Growth Characteristics

The growth pattern for MhCL03-F, MhCA04-F, and MhCB05-F CAFs was studied by using Alamar blue assay. Cells were seeded at a density of 2×10^3^ cells per well in a 96-well plate, and incubated at 37□ for 24 h. 12μL of Alamar blue solution (cat. no. DAL1025; Invitrogen; ThermoFisher Scientific, Inc.) was added resulting in a final concentration of 10% followed by an incubation for 18 h at 37□. The fluorescence readings were taken by collecting the supernatant media from the wells in Biotek Synergy H1 Hybrid Multi-Mode reader at 560nm and 590nm as excitation and emission wavelength respectively. The results display the mean ± standard deviation of three independent experiments.

### Immunocytochemistry

For detection of HPV by immunocytochemistry, respective cell lines at a concentration of 5×10^3^ cells per coverslip were cultured for 24 h followed by staining with p16 antibody (cat. no. G175-405; Biogeneics; Proteogen) (22). HeLa cells were used as a positive control for the experiment. Cells were fixed with 4% paraformaldehyde (cat. no. GRM3660-500g; HiMEDIA) for 10 min, followed by permeabilization with 0.1% Triton X 100 (cat.no. 10655; Fisher Scientific; Invitrogen) for 10 min at room temperature. Cells were blocked for 1 h at room temperature with 1% BSA (cat. no. TC194; HiMEDIA laboratories; LLC) in 1X PBS, followed by probing with p16 antibody and incubation at room temperature for 2 h. Secondary antibody (cat. no. K5007; Dako; Agilent Technologies, Inc.) was added and incubated for 30 min in dark. Visualization was done using 3,3′-Diaminobenzidine, and counterstaining with haematoxylin stain (cat.no. S034; HiMEDIA Laboratories, LLC). Mounting was done using DPX mounting media (cat. no. DAL1025; Qualigen Fine Chemicals TM; ThermoFisher Scientific) and observed under Nikon Eclipse E200 light microscope. The coverslips were washed twice with 1X PBS after every treatment.

### HPV PCR

PCR was run to check for the presence of HPV infections in the established CAF cell lines. The primers used were MY09 (5’CGTCCMARRGGAWACTGATC3’) and MY11 (5’GCMCAGGGWCATAAYAATGG3’) which flanks a 450 bp amplicon. As a control for the reaction, the DNA samples were subjected to PCR to amplify β actin using 5’ AGCCATGTACGTTGCTATCCA-3’ (Forward primer) and 5’ ACCGGAGTCCATCACGATG-3’ (Reverse primer) which flanks 120 bp amplicon was the reference gene control. Genomic DNA from HeLa cells was used as a positive control. PCR was set up using template DNA (200 ng), Taq polymerase (cat. no. D1806; Sigma-Aldrich; Merck KGaA), 1X PCR buffer, dNTPs (0.2 mM each), and primers (0.1 μM each). The amplification conditions for PCR were: 94□ for 5 min, 95□ for 30 sec (denaturation), 50□ and 60□ for MY09/MY11 and β actin, respectively, for 1 min (annealing) and 72□ for 1 min (elongation). Final elongation step was carried out at 72□ for 10 min. Visualization of amplified PCR products was done via agarose gel electrophoresis (1.5%).

### Flow cytometry

1×10^6^ cells/100μl were twice washed in PBS, followed by permeabilization with 0.1% triton X 100 (cat. no. 10655; ThermoFisher Scientific, Inc.) for half an hour followed by incubation with primary antibodies for 1 h on ice. Anti-FSP1 (cat. no. F4771; Sigma-Aldrich; Merck KGaA), EpCAM (cat. no. 4545; Cell Signalling Technology; Inc.), CD31 (cat. no. 303101; Biolegend; Inc.), and CD45 (cat. no. 361901; Biolegend; Inc.) were used as primary antibodies as per the dilutions instructed by the manufacturer. The cells were then treated with Alexa-488 conjugated secondary antibody (cat. no. A11029; Invitrogen; Thermo Fisher Scientific; Inc.) for 30 min in dark at room temperature. The cells were washed twice with 1X PBS after every treatment followed by centrifugation at 500 x g for 5 minutes at 4□. Unstained cells were used to correct the background fluorescence. The BD FACS Canto II system was used to conduct the experiment in duplicates, and the BDFACS Diva software, version 8.0.1, was used to analyse the results.

### Immunofluorescence

Purity of the established CAF cell lines were determined by staining 2×10^4^ cells, seeded on a coverslip and incubated overnight, fixed with 4% paraformaldehyde (cat. no. GRM3660-500g; HiMEDIA) for 10 min at room temperature, followed by permeabilization with 0.2% triton-X 100 in 1X PBS, blocked with 1% BSA (cat. no. TC194; HiMEDIA laboratories; LLC) in 1X PBS for 1 h at room temperature, followed by staining with anti-FSP-1,EpCAM, CD31, and CD45 antibodies and incubated for 2 h at room temperature. Cells were incubated with Alexa-488 conjugated secondary antibody for 1 h in dark at room temperature. DAPI histology mount (cat. no. F6057; HiMEDIA laboratories; LLC) was used to mount the coverslips on the slides. The slides were visualized under Zeiss Scope A1 fluorescent microscope using FITC filter. The coverslips were washed with 1X PBS after every treatment.

### STR Profiling

In order to determine the genomic identity and to exclude any cross-contamination of cell lines, STR profiling was performed using 10 loci as standard markers. Briefly, the genomic DNA was isolated from 1 × 10^6^ cells from all the cell lines. 50 ng of genomic DNA was used for the profiling. STR multiplex assay was performed by TheraCUES, Bangalore using GenePrint 10 (Promega corporation), version 3.0.0 of the SoftGenetics. GeneMarker_HID was used in order to analyze the result and the data was examined by referring to the STR Database of ATCC and CLASTR.

### DNA ploidy determination

Ploidy analysis was used to determine the cell lines’ DNA content (21). Normal cells from a healthy donor were chosen as a diploid control for the experiment because they rarely show any altered DNA content (22) and because their average DNA value has been classified as diploid (23). The ploidy of the cells was anticipated by determining the DNA index of the cells. The DNA was stained by treating the cells in PBS containing RNase (10 μg/ml; cat. no.12091021; Invitrogen; ThermoFisher Scientific, Inc.) and Propidium Iodide (40 μg/ml; cat.no. P4170; Sigma-Aldrich; Merck KGaA) at 37□ for 30 min. DNA content was determined in comparison to PBMCs which were used as control (diploid gDNA content). The fluorescence was determined by using BD FACS Canto II system to analyze the cells. The DNA index of the cell lines was estimated by dividing the mean channel of G0 phase cells by the mean channel of the lymphocytes.

### Isolation of human lymphocytes

Lymphocytes were isolated from a 32 year healthy female from Rajasthan, India, in December 2022 by collecting whole blood (diluted 1:3 in PBS) and layering above Ficoll Histopaque (cat. no. LSM-1077; HiMEDIA Laboratories; LLC) at 1:1 volume/volume ratio followed by centrifugation at 400 x g for 25 min at 20□. The acceleration and deceleration were maintained at 5 and 6 respectively throughout the procedure. The resultant buffy coat having lymphocytes were further washed with 1X PBS for further processing.

### Collection of Conditioned medium

CAFs were cultured in 6 well plates (cat. no. 140675; Thermo Fisher Scientific; Inc.) in complete RPMI until 80% confluency followed by changing the culture medium to serum free. Conditioned medium (CM) was collected after 48 h. of culturing the cells in complete RPMI medium without FBS, supplemented with 1X PenStrep and 1X GlutaMax.

### Estimation of proliferation

Epithelial cells (MhCT08-E and MhCT12-E), 1×10^4^ cells per well in a 96-well plate were treated with 50% serum free CAF condition medium supplemented with 10% FBS (cat. no. 10270-106; Gibco) and 50% fresh RPMI medium for 72 h. Cell viability was measured with Alamar blue as described for Growth pattern measurement above. Fold proliferation was estimated in comparison to without treatment with CAF conditioned medium. The results display the mean ± standard deviation of three separate experiments.

### Invasion assay

For invasion assay, ECM gel (cat. no. E1270; Sigma-Aldrich; Merck KGaA) was prepared in RPMI serum-free media at a final concentration of 1mg/mL. Cell culture inserts with a pore size of 8.0 μm (cat. no. TCP083; HiMEDIA Laboratories, LLC) were were coated with 100 μl of ECM and incubating at 37□ for 3-4 h, followed by seeding the cells on the apical chamber (over the ECM gel) with a density of 1×10^4^ epithelial cells per insert. Neat CAF Condition media was added in the basolateral chamber. The cells were allowed to invade for a period of 48 h and prepared for imaging by separating the cells on the inside of the insert using a cotton swab. Fixing of the cells was done by using 4% paraformaldehyde at room temperature for 10 min. The cells were then stained with 2% crystal violet for 5 min at room temperature. Any unbound dye was removed by washing it thrice with 1X PBS. The cells were then air-dried and imaged. The inserts were incubated in 10% acetic acid (cat. no. Q21057; Qualigen; Thermo Fisher Scientific, Inc.) in water while shaking at room temperature for 10 min to extract bound crystal violet for invasion quantitation. Absorbance was read at 590nm by transferring the extract into a 96-well microplate. The results display the mean ± standard deviation of three independent experiments.

### Sphere formation assay

A total of 5×10^3^ cells were seeded in the ultra-low (cat. no. 3473; Corning; Inc.) attachment plate along with the CAF conditioned medium. The cells were observed at 5^th^ day for their sphere formation ability and images were captured using bright field Nikon microscope. Size of the spheres was calculated using the formula 4/3πr^3^, where ‘r’ is the geometric mean of the two longest diameters of the spheres as calculated using Fiji ImageJ software vs. 1.53t.

### Statistical analysis

The mean and standard error of the mean are used to express all quantitative data. Unless otherwise noted, statistical significance was determined using the paired Student’s t-test. P < 0.05 was taken into consideration to show a statistically significant difference. All the statistical calculations were performed using GraphPad Prism software (version 5.00; GraphPad Software, Inc.).

## Results

### Characterization of established cell lines

#### Growth Characteristics

The three cell lines revealed a similar growth pattern as shown in Fig. 1A, wherein steady growth was observed until Day 5. However, MhCB05-F exhibited a more profound growth followed by MhCA04-F and MhCL03-F.

**Figure 1.**
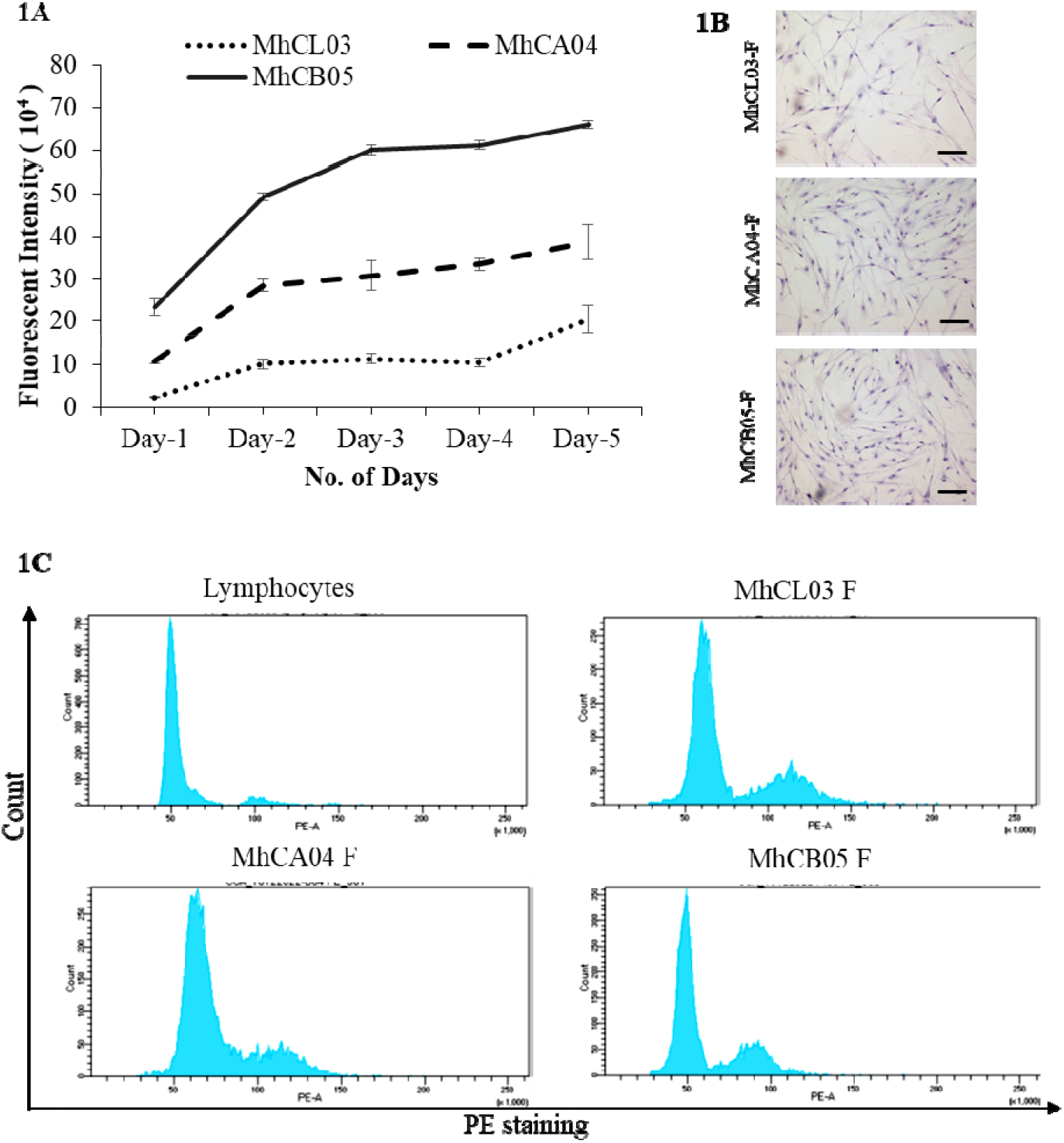
Characterization of established cell lines. (A) Growth pattern of MhCL03, MhCA04, and MhCB05 cell lines. (B) Microscopic images of Hematoxylin stained cells exhibiting spindle shaped morphology (Magnification 4X, Scale bar-100μm). (C) Determination of ploidy by flow cytometry analysis. Human lymphocytes, MhCL03, MhCA04, and MhCB05 stained with propidium iodide.

#### Morphology

The morphology of the established CAF cells was determined by staining them with haematoxylin and further observing under light microscope. As shown in the Fig. 1B, the cells exhibited a typical elongated, spindle shaped morphology (23,24), consistent with the fibroblast morphology as reported earlier (20).

#### STR profiling

In order to indicate the distinctiveness of the cell lines from those that are present in the ATCC and CLASTR database, STR profiling was done. As depicted in table III, upon analysis it was observed that none of the cell lines matched with the established cell line databases, as well as they were distinct from each other, indicating their novelty and excluding any cross contamination.

**Table III.** Proof of novelty of established cell lines by STR (<55% match cut-off from ATCC, CLASTR public databases, and among themselves)

|  |  | Parental | Passage /<br>Daughter | Parental | Passage /<br>Daughter | %<br>Match |
| --- | --- | --- | --- | --- | --- | --- |
| Sample Name | Marker | Allele#1 | Allele#1 | Allele#2 | Allele#2 |  |
| MhCL03-F & MhCA04-F | TH01 | 9 | 7 | 9 | 7 | 40.0 |
|  | D21S11 | 31 | 30 | 31.2 | 31.2 |  |
|  | D5S818 | 11 | 11 | 13 | 11 |  |
|  | D13S317 | 8 | 8 | 8 | 10 |  |
|  | D7S820 | 8 | 10 | 11 | 12 |  |
|  | D16S539 | 9 | 9 | 10 | 11 |  |
|  | CSF1PO | 11 | 11 | 12 | 13 |  |
|  | AMEL | X | X | Y | Y |  |
|  | vWA | 17 | 15 | 18 | 17 |  |
|  | TPOX | 8 | 8 | 10 | 9 |  |
| MhCA04-F & MhCB05-F | TH01 | 7 | 9 | 7 | - | 42.11 |
|  | D21S11 | 30 | 28 | 31.2 | 32.2 |  |
|  | D5S818 | 11 | 11 | 11 | 11 |  |
|  | D13S317 | 8 | 8 | 10 | 12 |  |
|  | D7S820 | 10 | 8 | 12 | 11 |  |
|  | D16S539 | 9 | 13 | 11 | 13 |  |
|  | CSF1PO | 11 | 10 | 13 | 12 |  |
|  | AMEL | X | X | Y | Y |  |
|  | vWA | 15 | 15 | 17 | 18 |  |
|  | TPOX | 8 | 8 | 9 | 9 |  |
| MhCL03-F & MhCB05-F | TH01 | 9 | 9 | 9 | - | 52.6 |
|  | D21S11 | 31 | 28 | 31.2 | 32.2 |  |
|  | D5S818 | 11 | 11 | 13 | 11 |  |
|  | D13S317 | 8 | 8 | 8 | 12 |  |
|  | D7S820 | 8 | 8 | 11 | 11 |  |
|  | D16S539 | 9 | 13 | 10 | 13 |  |
|  | CSF1PO | 11 | 10 | 12 | 12 |  |
|  | AMEL | X | X | Y | Y |  |
|  | vWA | 17 | 15 | 18 | 18 |  |
|  | TPOX | 8 | 8 | 10 | 9 |  |

ATCC: American Type Culture Collection; CLASTR: Cellosarus STR Similarity Search Tool; E: Epithelial; F: Fibroblast

#### Ploidy determination

DNA index was calculated as the ratio of the mean fluorescent intensity of the CAFs and healthy diploid lymphocytes in G0/G1 phase. Upon analysis it was observed that MhCL03-F, MhCA04-F and MhCB05-F cells had DNA indices of 1.2, 1.3, and 0.94 respectively (Fig. 1C). The findings suggested that the patient samples contain aberrant DNA, which may be the cause of these cells’ immortalization.

#### Purity

In order to use a cell line as a study model for in vitro experimental studies, it is important to authenticate and confirm the purity of the cell lines established. The origin of cancer-associated fibroblast was determined by confirming the expression of fibroblast-specific markers via flow cytometric analysis. As shown in Fig. 2A, the CAF nature of the cell lines was confirmed by positive staining with Fibroblast specific protein – 1 (FSP-1) antibody. Similarly, the origin of CAFs was also confirmed by negative staining with EpCAM antibody which is specific against epithelial cells (25). FACS analysis results revealed that, MhCL03-F, MhCA04-F, and MhCB05-F exhibited a percentage positivity of 99.7%, 95.2%, and 89.4% for FSP-1 respectively. Whereas, observed for EpCAM expression. The purity of cell lines was < 0.1% positivity was also determined via immunofluorescent analysis (Fig. 2B). The cells showed positive staining for FSP-1 antibody which is specific against fibroblast cells. Apart from this, negative staining of the cells against EpCAM, CD31, and CD45 antibodies which are specific to epithelial cells, endothelial cells and leucocytes respectively, confirmed the CAF nature of the established cell lines (26). Similarly, the cells stained negative for CD31 and CD45 antibodies in flow analysis as well (<0.1% population positivity), keeping human lymphocytes as the control, further suggesting that established cultures are from fibroblast lineage only (Fig. 2C). MhCT12-E and MhCT08-E epithelial cells which were previously established and characterized by Dwivedi et. al (20) were used as a neg tive control for FSP-1 staining and positive control for EpCAM staining. Overall, the results conclude and confirms the purity and origin of established CAF cell lines.

**Figure 2.**
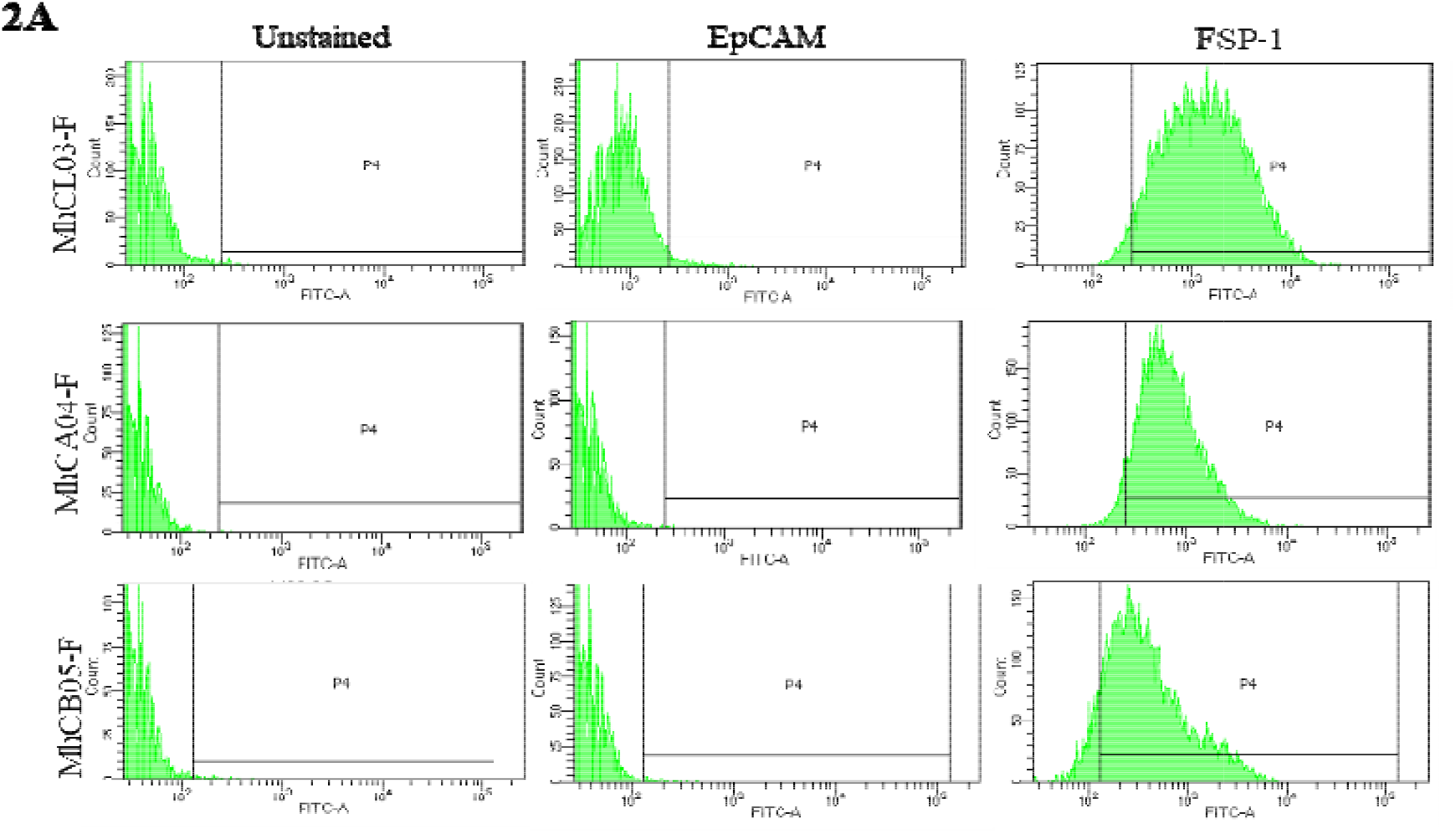
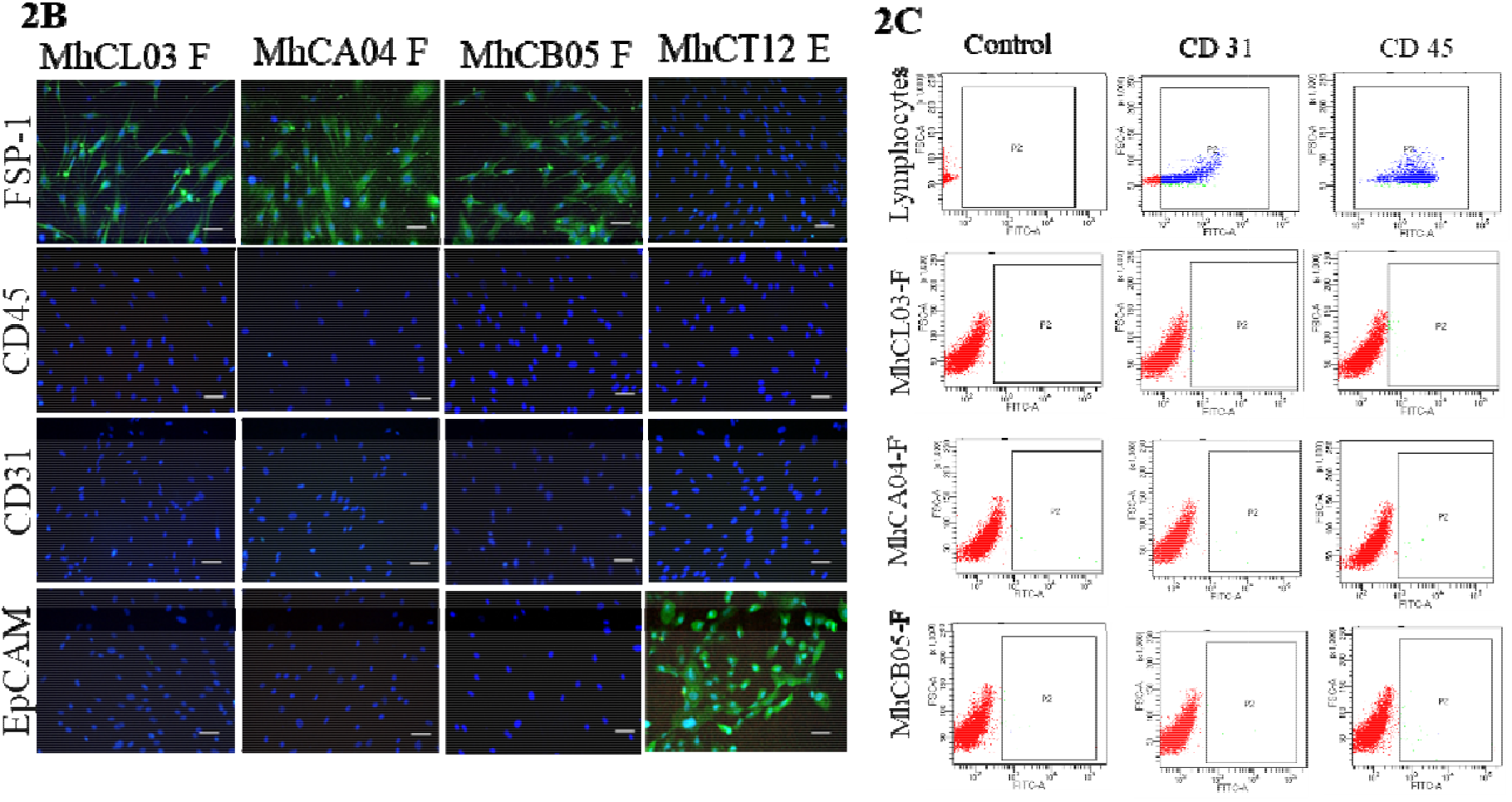
Purity of established cell lines. (A) FACS analysis displaying EpCAM and FSP-1 staining for the established cell lines. (B) Fluorescent images of established cell lines stained with F P-1, EpCAM, CD31, and CD45 (Magnification 10X, Scale bar-50μm). (C) FACS analysis displaying CD31 and CD45 staining for the lymphocytes and the established lines as indicated.

#### HPV detection

HPV infections are recognized as one of the major factors leading to HNSCC. It has been studied that, HPV positive HNSCCs exhibits a much convenient prognosis when compared to HPV negative cancers (27). Thus, it becomes important to determine the HPV infection status of an established cell line. Immunocytochemistry analysis was carried out to determine the status of HPV infection, wherein expression of p16 antigen was examined (22). Upon PCR and immunocytochemical analysis, it was observed that, all the three cell lines were stained negative for p16 antibody (Fig. S1A) as no nuclear stain with the p16 antigen was observed in any of the established cultures, while the HeLa culture stained positively with the same. HPV negative status of the established cell lines was further confirmed by PCR as no amplified bands were observed (Fig. S1B), however, a 120bp amplicon was observed in beta actin control.

#### Tumorigenic properties of established cell lines

Several assays were carried out to determine the tumorigenic potential of the CAFs, by treating the epithelial cells with CAF condition medium

#### Estimation of proliferation

Upon treatment with CAF conditioned medium, a significant increase in proliferation of epithelial cells was observed (P<0.01) as compared to the no treatment control, wherein maximum significant proliferation was seen at 48 h for all the five cell lines (Fig. 3A). Upon comparison between all three cell lines and previously developed two tongue-CAFs (Fig. 3A), it was observed that the MhCA04-F conditioned medium had the best and most significant effect on the induction of proliferation in both the MhCT12-E and MhCT08-E cell MhCB05-F conditioned medium followed next, while MhCT08-F and lines. MhCL03-F and MhCT12-F conditioned medium were similar to MhCB05. Interestingly, MhCT08-E cell line showed higher proliferation after 24 h of treatment with CAF conditioned medium as compared to MhCT12-E cell line.

**Figure 3.**
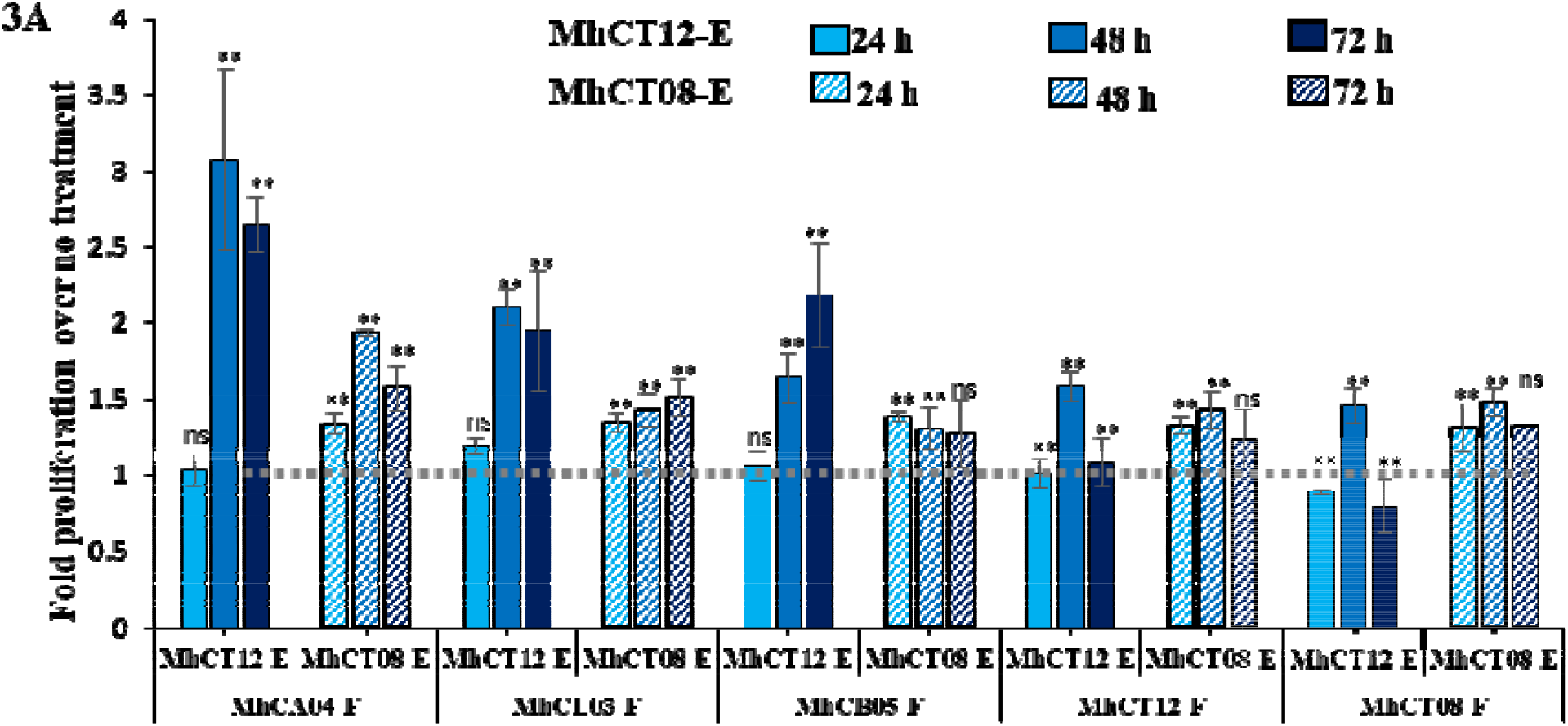
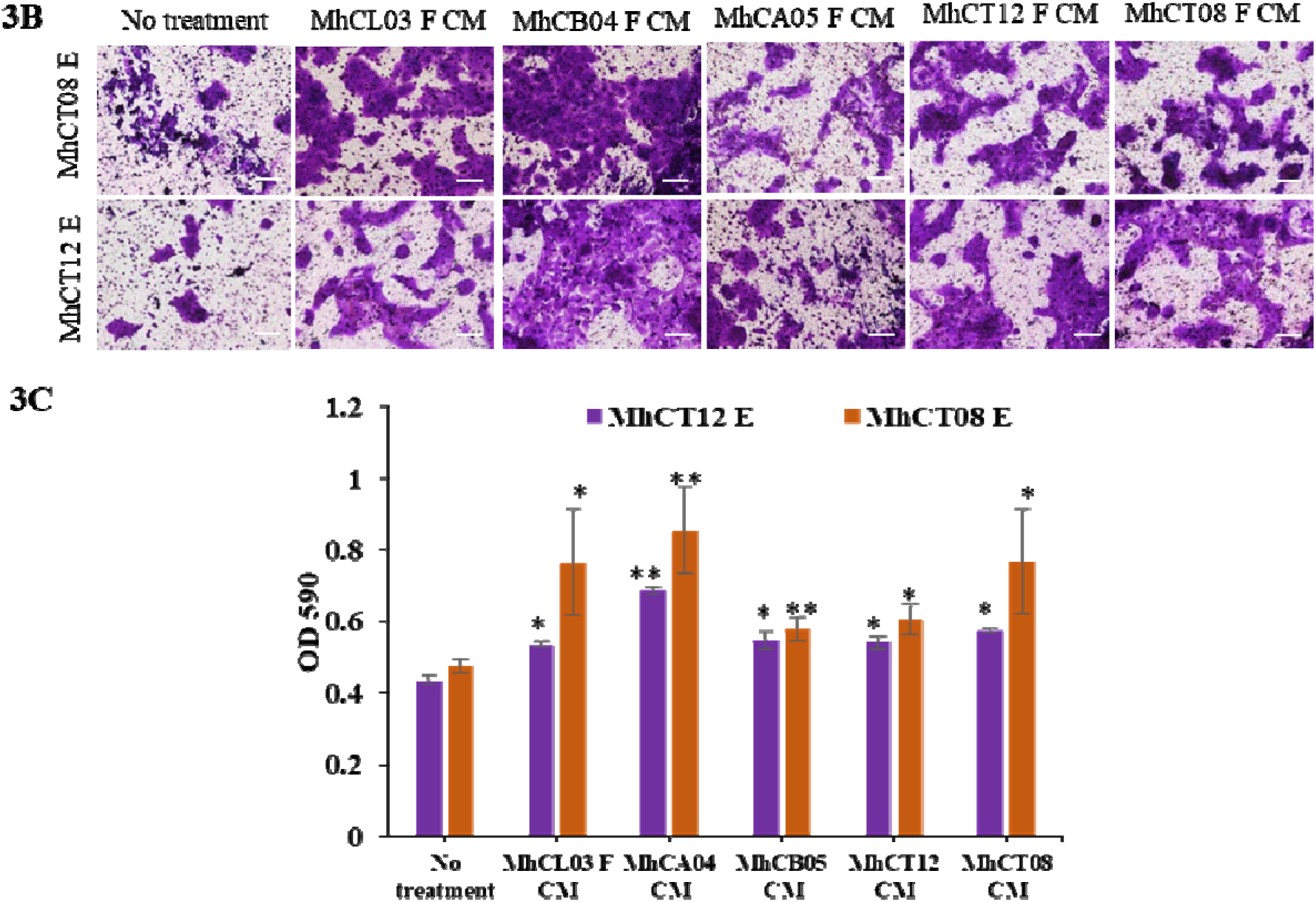
Tumorigenic properties of established CAF cell lines. (A) Proliferative potential of MhCT12-E and MhCT08-E under the influence of indicated conditioned medium. The grey line intersection depicts the no treatment control for both the epithelial cells. Statistical significance over no treatment, where, *p<0.05 and **p<0.01 (B) Invasive potential of epithelial cells under the influence of CAF condition medium (Magnification 10X, Scale bar-100μm). (C) Graphical illustration of the invasive effect of CAFS on epithelial cells. CM-Conditioned medium, E-Epithelial, F-Fibroblast.

However, proliferation rate of MhCT12-E cell line increased dramatically as compared to MhCT08-E cell line upon prolonged treatment with indicated conditioned medium. This was due to the inherently increased proliferative ability of MhCT08-E cells as compared to MhCT12-E cells. Upon prolonged culture for 48-72 h, MhCT08-E cells proliferated faster even without any treatment with CAF conditioned medium (Figure S2), therefore, upon normalizing with the no treatment group, the overall effect of CAF conditioned medium was not as high in MhCT08-E cells when compared to the MhCT12-E cells.

#### Invasion assay

A significant increase in the invasive potential of MhCT12-E and MhCT08-E epithelial cells was observed upon treatment with CAF condition medium (Fig. 3B,C). Upon quantification, it was observed that the maximum invasion by both the epithelial cells was observed under the influence of MhCA04-F condition medium (P< 0.01) followed by MhCL03-F condition medium (P<0.05) and MhCB05-F conditioned medium as compared to the no treatment control. Previously reported CAF conditioned medium also increased the invasive ability of the cells (Fig. 3B,C), however, not as much as MhCA04-F cells.

#### Sphere formation

Sphere formation potential of epithelial cells significantly increased under the influence of CAF condition medium (Fig. 4). Upon analysis it was observed that not only the size (Fig. 4A), but also the total number (Fig. 4B) of the spheres was significantly higher upon treatment with CAF conditioned medium as compared to the no treatment group. Similar to the other tumorigenicity assays, the effect of the MhCA04-F CAF conditioned medium was the most profound followed by MhCL03-F, MhCA05 CAF, MhCT08 CAF and MhCT12 CAF conditioned medium (Fig.4C). Upon quantification, it was observed that the total number of spheres formed upon treatment with MhCL03-F, MhCA04-F, and MhCB05-F CAF condition medium were 40, 93 and 42 in case of MhCT08-E cells (Fig. 4B). Similarly, treatment with MhCL03-F, MhCA04-F and MhCB05-F CAF condition medium resulted in a total of 19, 29, 17 spheres respectively in MhCT12-E cells (Fig. 4B). While treatment with previously established MhCT12-F and MhCT08-F CAF condition medium resulted in 21 and 36 spheres in MhCT12 Epithelial cells, while 57 and 66 spheres in case of MhCT08 Epithelial cells. Additionally, as per the previous report, it was also observed that MhCT08-E formed much larger and number of spheres under the influence of CAF condition medium when compared to MhCT12-E cells (Fig.4 C).

**Figure 4.**
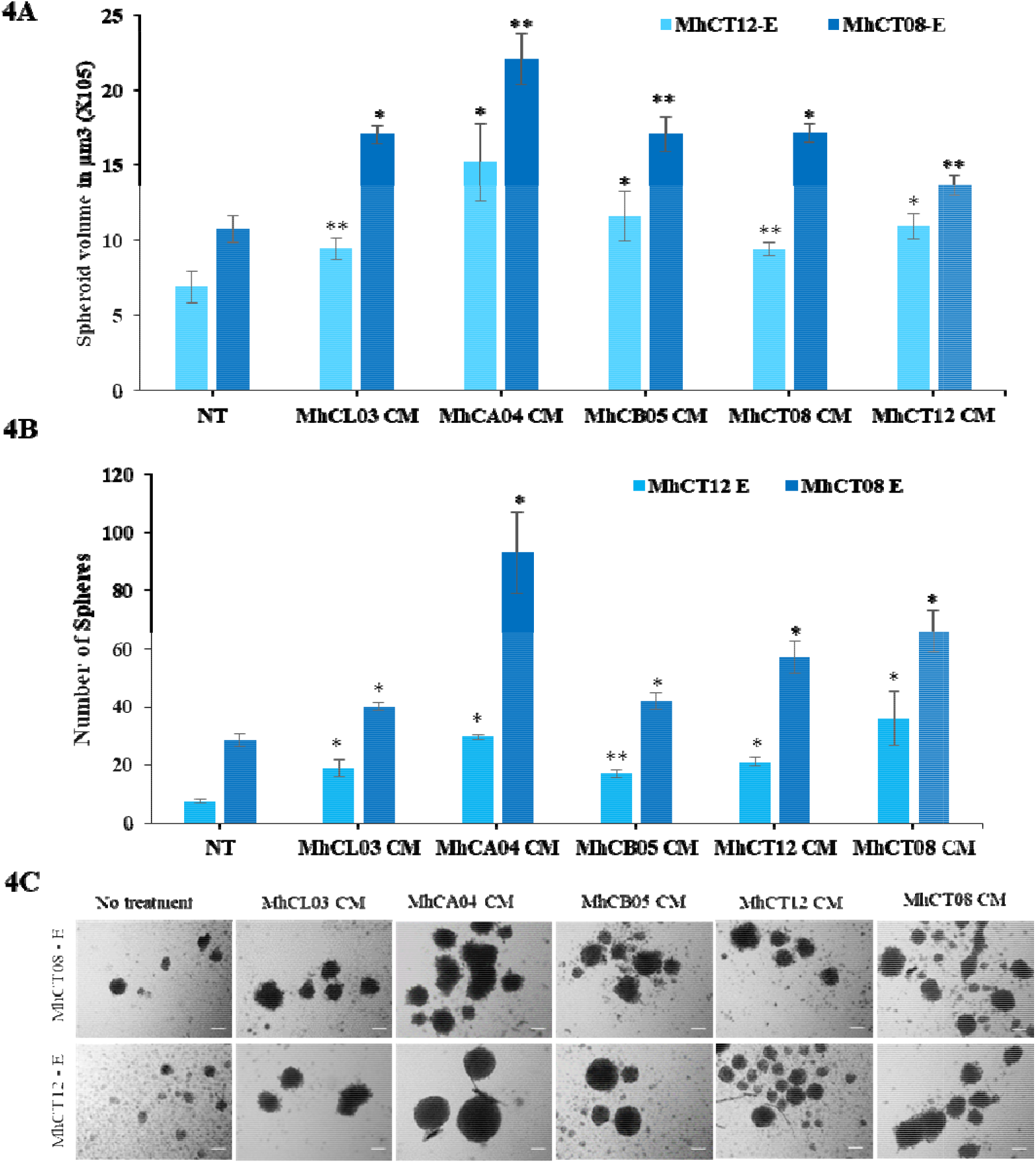
Sphere formation assay. (A) Volume of MhCT08-E and MhCT12-E spheres formed under the effect of CAF conditioned mediums as indicated (*p<0.1, **p<0.05), (B) Sphere forming potential of MhCT08-E and MhCT12-E cells under the influence of CAF condition medium as indicated (*p<0.1 and **p<0.01), Statistical significance indicates difference between no treatment and CAF conditioned medium treatment (C) Sphere formation potential of MhCT08-E and MhCT12-E cells under the influence of CAF condition medium. (Magnification 10X, Scale bar-100μm), CM-Condition media, NT-No Treatment, F – Fibroblast, E – Epithelial.

## Discussion

Over the past decade it has become evident that TME encourages tumor growth (28). Concern has been raised about the ECM’s constituent parts and mechanical attributes as significant catalysts for the behavior of cancer cells and the development of the disease. CAFs are a crucial component of TMEs and are essential for matrix remodeling (29). CAFs are responsible for the secretion of multiple growth factors, kinases, cytokines, and chemokines into the TME to facilitate tumor progression (30–33). Thus establishment of CAF cell lines from HNSCC will be of immense value to the scientific community and eventually to the clinic through translational research.

In this study along with the previous one (20), we report a total of 5 CAFs from 4 sites of lesions of 5 HNSCC patients.

Development of cell lines from head and neck cancer patients from India has greatly advanced the understanding of tumor heterogeneity and revealed the molecular variations between the Indian (Asian) and Western populations. Populations from Western nations and Southeast Asian regions have a vast range in HNSCC epidemiology, which may be caused by genetic variations between these populations (34). A source of cancer cells with the heterogeneity of the cell populations present in the parental tumor may be found in the cell lines developed from patients with HNSCC. Notably, they can be utilized to comprehend the molecular processes underlying cancer cells’ resistance to chemotherapy and radiation treatment (35). In the current investigation, three Indian males with known risk factors diagnosed with squamous cell carcinoma of the larynx, buccal mucosa, and upper alveolus were used to produce and characterize three unique CAF cultures.

Primary cultures have been developed using a variety of techniques. Explant culture is a method that has been used to create cell lines in numerous investigations (35–39). Several studies also mention seeding the cell suspension and directly digesting the tumor tissue (17,40), Using a mouse xenograft model, Mulherkar et al. (41) revealed the creation of NT8e, an oral squamous cell cancer cell line. Explant culture was used in the current investigation. According to a recent study, different media can be used to promote the populations of fibroblasts and epithelial cells to develop at different rates (42). To avoid any phenotypic or genotypic changes brought on by patient-derived xenograft generations, the cultures described in the present investigation were grown in RPMI-1640 complete media as also in the previous report (20). These cultures did not require any feeder cells or viral vectors to be stabilized spontaneously. The cell lines stained positive for FSP-1 via both the immunocytochemical and flow cytometric analysis. Additionally, while endothelial cells and fibroblast cells have a very similar appearance, negative staining with the endothelium specific (CD31) and hematopoietic specific (CD45) markers disproved their suspicion of cross contamination in the established cultures and proving their fibroblast specific lineage.

An insight of the aggressiveness, potential for metastasis, and prognosis of the disease can be gained through DNA ploidy analysis of cancer cells (43–45). The three cell lines contained aneuploid DNA than the lymphocytes’ diploid DNA. While the DNA indices of MhCB05-F (0.94) indicated aneuploidy, DNA indices of MhCL03-F (1.2) and MhCL04-F (1.3) implied hyperploid DNA content. The quick and frequent cell divisions of cancer cells may explain this situation. Uncontrolled cell divisions lead to aberrant karyokinesis, which raises the ploidy level of the cells as a result of improper karyokinesis (46).

One of the main risk factors for the emergence of HNSCC is HPV infection. According to the literature, HPV infection raises the risk of mouth cancer (47). By employing MY09/MY11 primers for the virus’ capsid protein, PCR was used to test the current cell lines for the presence of HPV markers. HPV infection status was also checked by looking for nuclear stain of p16 antigen. The current findings showed that no HPV infection was present in any of the three cell lines. As demonstrated previously, even though these epithelial cell types had oncogenic potential, treatment with the three fibroblasts’ conditioned medium significantly increased the proliferative, invasive and sphere formation ability of both the epithelial cells thus demonstrating that all the five CAFs possessed the capability to elevate the oncogenic potential of the tumor epithelia. In this study we observed that even though the CAFs were from different head and neck sites, upon co-culture, they were still able to increase the tumorigenicity of the both the epithelial cells derived from tongue squamous cell carcinoma, a different site altogether. This implies that these CAFs are not restricted in imparting the increased tumorigenicity to a particular site, but may also be used to study tumor progression from a number of sites in oral cancer, or may even be expanded to pan cancer studies, making them an immensely useful tool. This may be due to various signaling factors released by the CAFs, which promote tumorigenesis and establishes the autologous pair of cells as an effective model to study tumor□stroma crosstalk. Furthermore, upon analysis, it was observed that MhCT08-E cells were more tumorigenic as compared to MhCT12-E cells in vitro.

All the five CAFs under a co-culture setting with tumor epithelial cells help to simulate the actual interaction between tumor cells and the CAFs. Additionally, the CAFs themselves can serve as excellent model systems for studying tumor evolution. However, the study is limited to in vitro potential so far.

In conclusion, the cell lines generated in this work show consistent cell shape and enduring growth capacity. Along with the two autologous pairs established by the group earlier, they offer a practical tool for doing basic and translational research in the area of tumor-stroma interaction in head and neck cancer, in addition to having the potential to be employed as a novel drug testing platform in co-culture settings.

## Acknowledgements

The authors would also like to thank Dr. Anjali Karande from IISc for providing the HeLa cells.

## Availability of data and materials

All data generated or analysed during this study are included in this published article.

## Authors’ contributions

MD and AS conceived the study. MAK and VP provided the samples. CG collected the patient samples and isolated fibroblast cells. ND and SPK designed the characterization study. ND, HB, SD and CC performed the experiments. MD and ND wrote the manuscript, analysed and interpreted the data and confirm the authenticity of the manuscript. AS and SPK reviewed the manuscript. All authors read and approved the final manuscript.

## Ethics approval and consent to participate

The present study was approved [approval no. NHH/MEC-CL-2015-405 (A)] by the ethics committee of Narayana Health City (Bangalore, India). Informed consent for the study was obtained from all patients.

## Patient consent for publication

Not applicable.

