## Supplementary figures for "Establishment and Characterization of Three Novel CAF Cell Lines from HNSCC Patients"

### Slide 1
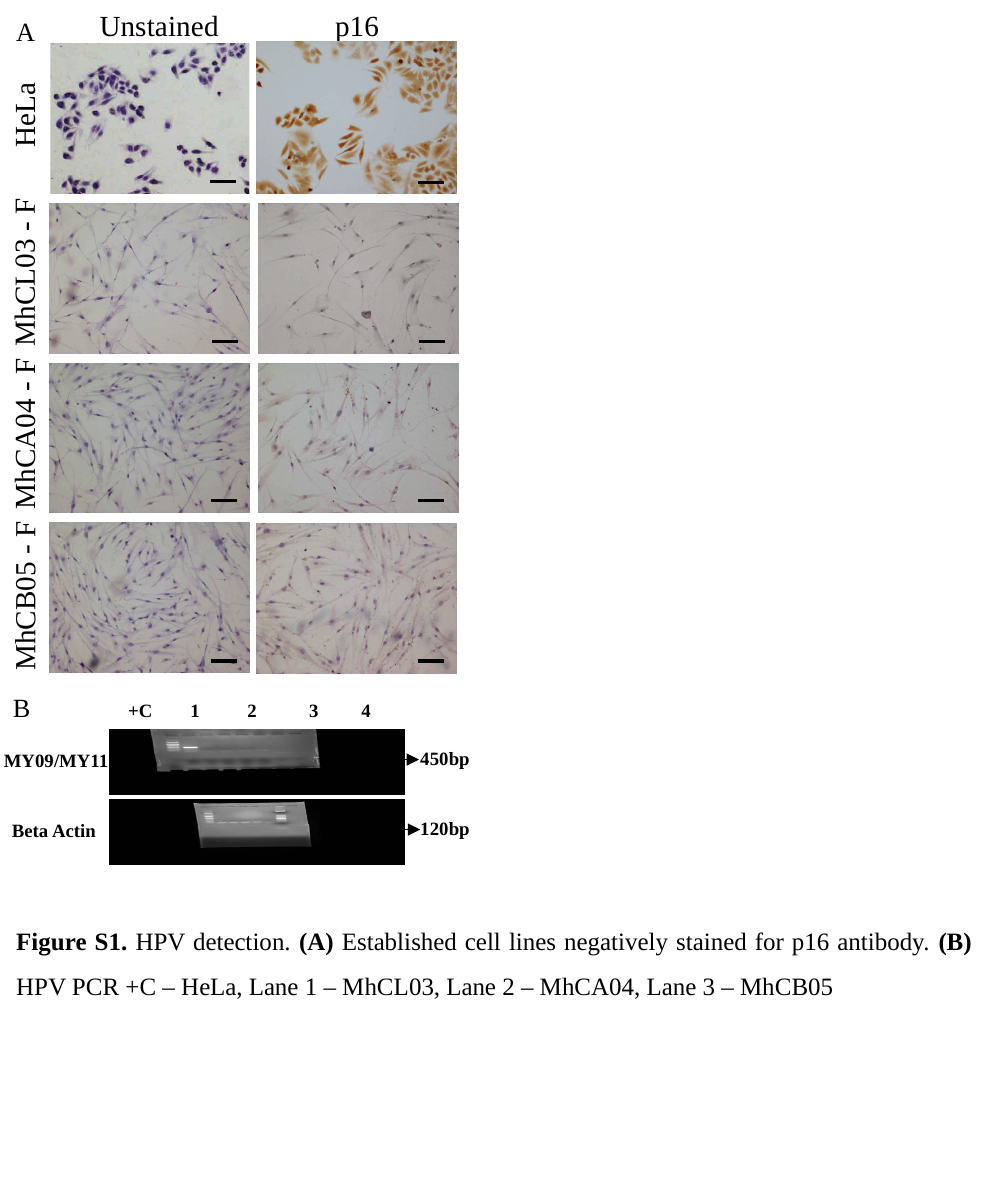

p16
Unstained
A
HeLa
MhCL03 - F
MhCA04 - F
MhCB05 - F
B
+C 1 2 3 4
450bp
MY09/MY11
120bp
Beta Actin
Figure S1. HPV detection. (A) Established cell lines negatively stained for p16 antibody. (B) HPV PCR +C – HeLa, Lane 1 – MhCL03, Lane 2 – MhCA04, Lane 3 – MhCB05

### Slide 2
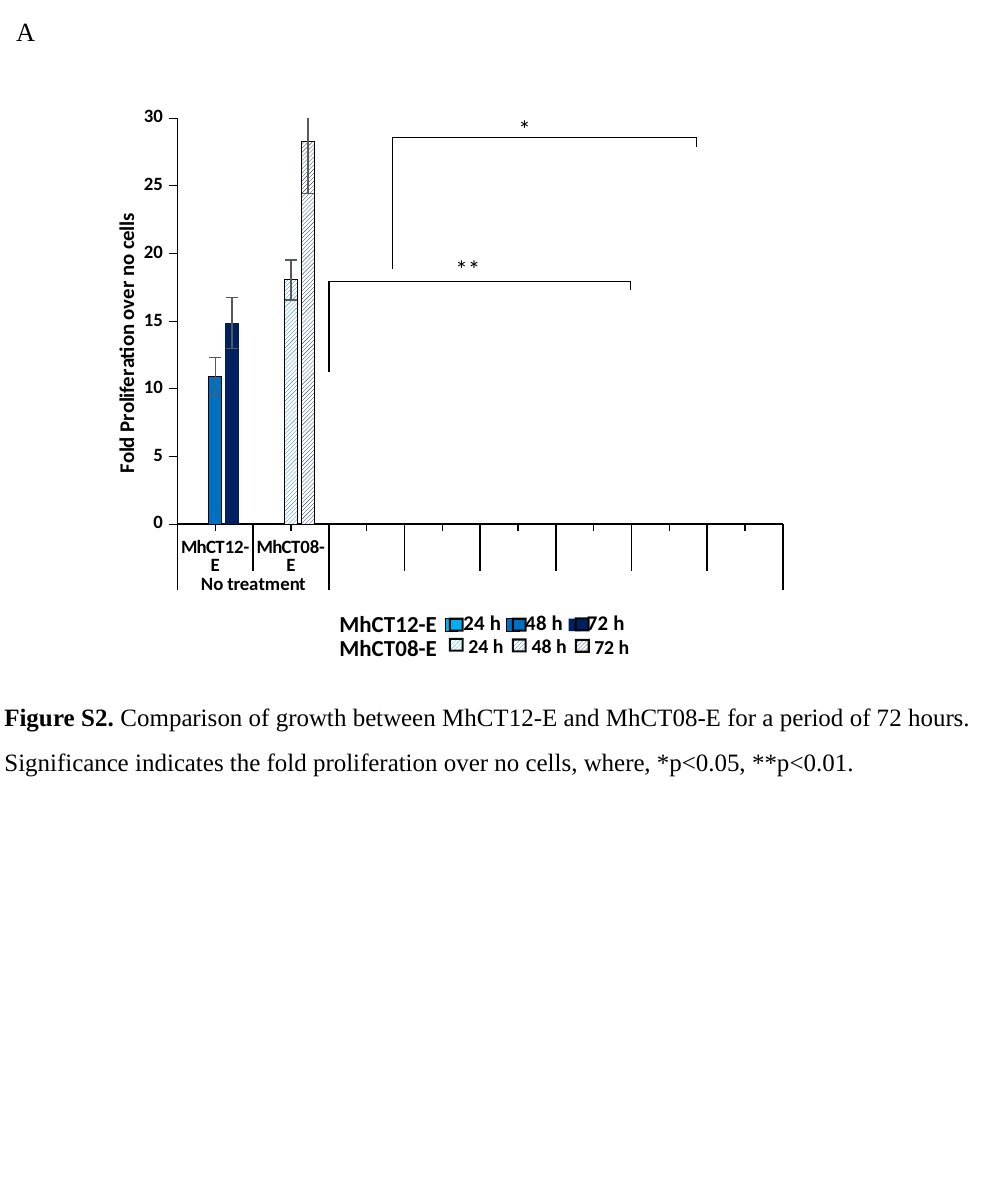

A
#### Chart
| Category | 24 h | 48 h | 72 h |
|---|---|---|---|
| MhCT12-E | 17.36852696806073 | 10.95807622132475 | 14.86780434331763 |
| MhCT08-E | 14.336069656697681 | 18.055729343083144 | 28.28990015702308 |MhCT12-E
MhCT08-E
24 h
48 h
72 h
Figure S2. Comparison of growth between MhCT12-E and MhCT08-E for a period of 72 hours. Significance indicates the fold proliferation over no cells, where, *p<0.05, **p<0.01.
